# Decoding spatial organization maps and context-specific landscapes of breast cancer and its microenvironment via high-resolution spatial transcriptomic analysis

**DOI:** 10.1101/2023.10.25.563904

**Authors:** Eun Seop Seo, Boram Lee, Inwoo Hwang, Ji-Yeon Kim, Kyeongmee Park, Woong-Yang Park

## Abstract

Single-cell RNA transcriptomics has revealed the intricate heterogeneity of both tumors and their microenvironment. However, a notable limitation is its inability to retain spatial context, a crucial aspect of understanding cell identity and function. In this study, we employed imaging-based single-cell spatial transcriptomics to elucidate the tumor and immunological landscapes of two breast cancer samples. By resolving over 400 000 cells per slide, we demonstrated that transcriptional differences lead to structural disparities within and between tumors. Additionally, we observed that the composition of the tumor microenvironment varies depending on its spatial location. Notably, we detected immune cell gradients transitioning from the tumor periphery to its core regions and from tertiary lymphoid structure to immune inflamed regions, in alignment with the specific function of each cell type. This finding facilitated a more precise classification of the tumor immune microenvironment. This study provides a comprehensive dataset for breast cancer researchers and underscores the significance of spatial context in understanding the multifaceted heterogeneity of cancer and its environment.

## Introduction

Recent advances in treatment strategies for breast cancer (BC) have significantly improved survival outcomes in patients.^1^ However, several patients diagnosed with advanced BC still have limited overall survival. Specifically, patients diagnosed with triple-negative breast cancer (TNBC) and human epidermal growth factor receptor-2 (HER2)-positive BC^2^ have a worse prognosis. Moreover, these patients show various outcomes despite being treated with the latest treatment regimens, including immune checkpoint inhibitors, antibody-drug conjugates, and target agents like trastuzumab and pertuzumab.^3–6^ Recent studies have revealed that the disparity in the outcomes of patients with BC can be attributed to tumor heterogeneity, including intratumoral and intertumoral heterogeneity.^7–9^ HER2 status has been reported to exhibit both intratumor and intertumor heterogeneity.^10,11^ Additionally, the intrinsic subtype determined by RNA-based multigene expression assays differed from the conventional BC subtype and had prognostic value within the same BC subtype.^12–14^

This heterogeneity in BC fundamentally stems from the fact that each tumor is composed of a complex arrangement of distinct regions and cells. While single-cell RNA sequencing (scRNA-seq) has identified various tumor cell subpopulations^15,16^, the tissue digestion method required to extract cells eliminates the spatial information. Nevertheless, recognizing the spatial relationships and interactions between these cancer cells and neighboring cells is pivotal, as it profoundly influences their behavior and inherent properties.^17,18^

Furthermore, comprehending the spatial landscape of the tumor immune microenvironment (TIME) is crucial for understanding BC. The specific positioning of immune and stromal cells within tumors reflects their dedicated function.^19,20^ Recent advancements in cancer immunotherapy for BC, such as checkpoint blockade, vaccines, engineered T cells, and NK cells, have generated interest in developing a universal approach to target TIMEs.^21^ However, the immune landscape varies across tissues and even between distinct regions of the sample tumor. Each site harbors a specific assortment of tissue-resident and systemic immune cells. Acknowledging the context-specific nuances of immune responses can guide the development of tailored immunotherapies for BC. Assessing the composition, proportion, and trajectory of TIME cells in a context-specific manner may help predict the efficacy of cancer immunotherapy.^22^

In this study, we investigated the spatially resolved features of BC using high-resolution *in situ* sequencing.^23,24^ We attempted to decode the spatial configurations, intrinsic heterogeneity, and tumor immune microenvironment in a context-specific manner. These data represent an initial step towards obtaining a comprehensive understanding of BC and can serve as a valuable resource for future studies.

## Materials and Methods

### Sample preparation

Two formalin-fixed, paraffin-embedded (FFPE) BC tissue blocks (one HER2-positive and one TNBC) that were banked after standard-of-care procedures were collected and subsequently processed in alignment with the manufacturer’s instructions, with approval from the Sanggye Paik Hospital Institutional Review Board. The molecular subtype of each sample was determined using routine immunohistochemistry (IHC) procedures. From each FFPE sample, sections of 5 µm thickness were carefully sectioned and placed on Xenium slides (10x Genomics, Pleasanton, CA, U.S.A), each measuring 2 × 1 cm. The 280 DNA probes targeting the genes included in the Xenium Human Breast Panel (Supplementary Table 1), 20 negative probes and 20 decoding control probes were used. DNA probes were flanked by two regions that hybridized independently to the target RNA and contained a gene-specific barcode sequence. After the probe’s ends were ligated, a circular DNA probe was created. This resulted in the generation of multiple copies of the gene-specific barcode as the rolling circle product (RCP). Following cleaning, the background fluorescence was diminished. Subsequently, the sections were positioned in an imaging cassette and loaded onto the Xenium Analyzer instrument.

### Image processing and Decoding on Xenium Analyzer

The Xenium Analyzer performs both analytical detection and imaging through cycles. The instrument automatically administers a series of actions: introducing fluorescently labeled probes designed to detect the linearly amplified DNA as RCPs, allowing for incubation, capturing images of the sample, and removing of the probes. During each cycle, fluorescent oligonucleotides are hybridized to the amplified barcodes, allowing the measurement of fluorescence intensity across four distinct Xenium color channels. Consequently, each optical signature is decoded into a transcript identity with their location in x, y, and z recorded. Between each cycle, cell segmentation occurs. DAPI images are used to infer cell boundaries using a machine learning algorithm. Then cells are assigned to cells based upon the segmentation results. Once the transcripts are assigned to cells, the Onboard Analysis software generates a gene by cell matrix, performs secondary analysis and generates Xenium Explorer file for downstream interpretation. More information can be found on the 10x Genomics Support Site.

### Data processing, differential gene expression analysis, and cluster annotation

Following Xenium’s standard analysis, we structured our data into an anndata format for further processing using Scanpy (v1.9.5)^25^. Gene expression count data were log-transformed and normalized. Neighborhood graphs were computed considering 50 principal components and 40 neighbors, followed by Leiden clustering for each sample. Global cell clusters were annotated using differentially expressed marker genes and differential gene expression (DGE) analyses. The significance of the DGE was assessed using the Wilcoxon test on log-normalized count data. We also conducted gene set enrichment analysis (GSEA) on cancer clusters within the TNBC sample to determine whether epithelial-mesenchymal transition (EMT) pathways were upregulated in the Cancer 2 cluster which has a mesenchymal-like morphology.

### Histopathologic annotation and differential composition by compartments

The images captured from the Xenium slides were reviewed and annotated by a pathologist using Adobe Illustrator 2023 software. The regions of interest were indicated for each slide. After delineating a region of interest (ROI) by drawing a polygon around histologically distinct features, the cell type composition was compared to each ROI.

### Spatially variable genes

We computed Moran’s I global spatial autocorrelation statistic^26^ to investigate the spatial distribution of molecular subtype markers in each tissue using Squidpy (v1.3.0)^26^. This statistic is characterized by the correlation in a signal at nearby locations.

### Supervised clustering for minor cell type

Given the limited number of genes (280) available in the Xenium panel, unsupervised clustering was not feasible for more detailed cell identification. Consequently, we relied on supervised clustering for further annotations of T cells and macrophages, guided by a literature review.^27–30^

Following the extraction of the cluster of differentiation CD4 T cells, CD8 T cells, and macrophages, we subdivided the dataset according to the cell type. For further cell identification, we used the following markers: *SELL* and (*TCF7* for naïve T cells; *CCR7* and *IL7R* for central memory (CM) T cells; *GZMA* and *GZMK* for effector memory T cells; and a combination of *LAG3*, *PDCD1, TIGIT*, and *HAVCR2* for exhausted T cells. In the CD4 T cell cluster, *FOXP3* and *CTLA4* were used for regulatory T cells, whereas CTLA4 was used as an exhaustion marker for CD8 T cells. In the macrophage cluster, we associated *ITGAX*, CD80, CD86, *CCL5*, and *CCR7* with M1-like macrophages, and *MRC1* and *CD163* to M2-like macrophages. Macrophages that did not exhibit both features were classified as non-activated M0-like macrophages. Subsequently, we assigned cell types based on dominant gene expression in each cell type. Using the prominence of these gene markers, we meticulously categorized cells based on their genetic signatures.

### Spatial trajectory analysis

To identify the spatial trajectory during lymphovascular invasion in the TNBC samples, we designated the Cancer 1 cluster as the root and computed the pseudo-space-time distance using stLearn (v.0.4.9)^31^. Subsequently, we identified the transition genes along the Cancer 1 to Cancer 2 meta-trajectory by employing a Spearman correlation rho cutoff of 0.1.

### Ligand-receptor (L-R) analysis

We created a grid on the slide of the HER2-positive sample, sectioning it into 150 × 150 segments to increase the computation speed for extremely large Xenium data. Subsequently, we performed an ligand-receptor (L-R) permutation test to determine regions of high L-R co-expression using stLearn in the within-spot mode. Permutation tests for the enrichment of L-R pairs between two cell types compared to the random null distribution were performed using connectomeDB^32^.

After adjusting the significance thresholds, we visualized the scores and p-values associated with the expression of the target LR pairs.

## Results

### Single-cell spatial transcriptome profiling of breast cancer

The workflow of the spatial transcriptomic analysis is described in detail in the Methods section and illustrated in Fig. 1a. To elucidate the cellular architecture of BC, we analyzed 655 782 single cells in the HER2 positive sample and 482 965 single cells in the TNBC sample. Through Leiden clustering and annotations of differentially expressed genes, we identified nine major cell types in the HER2-positive sample and eight major cell types in the TNBC sample (Supplementary Fig. 1a–d). The eight identified cell types included cancer cells, myoepithelial cells, four immune cell types (myeloid, T, B, and plasmablasts), and two stromal cell types (mesenchymal and endothelial cells). Necrotic areas were isolated as separate clusters. Notably, myoepithelial cells were not detected in the TNBC sample because of the absence of ductal carcinoma *in situ* (DCIS) and normal lobules in the specimen. Subsequently, we mapped each cell type onto the spatial dimensions (Fig. 1b, c). The spatial distribution of all cell types is closely aligned with manual annotations made by a pathologist (Fig. 1d, e). As observed in the hematoxylin and eosin (H&E) staining images, the HER2 positive sample exhibited a higher proportion of cancer cells, whereas the TNBC samples demonstrated a higher proportion of mesenchymal cells (Fig. 1f, g).

**Fig 1.**
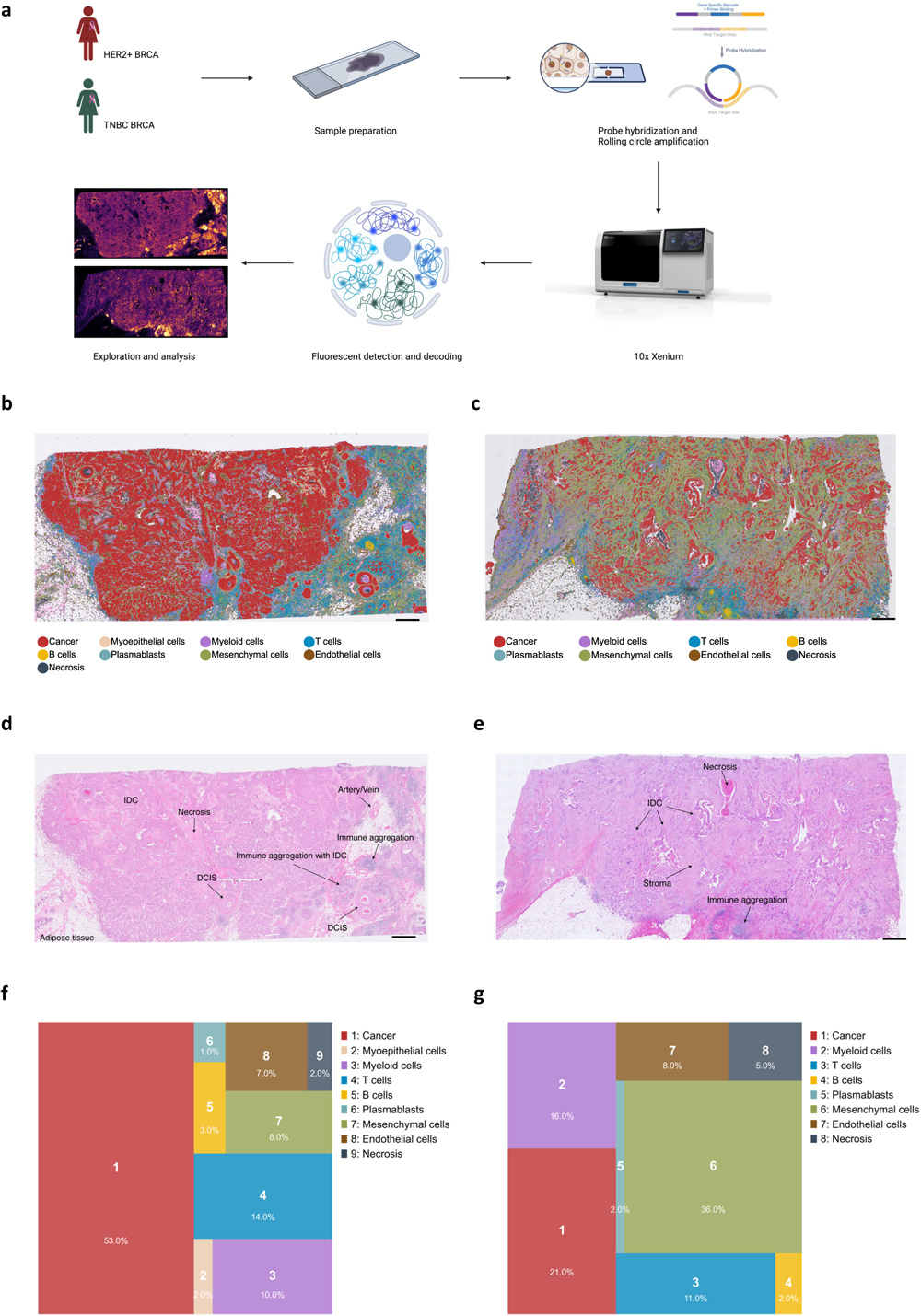
Study design and cell type mapping on space. a Overview of the study design. The image was created using BioRender.com. b, c Spatial mapping of the cell types in human epidermal growth factor receptor-2 (HER2)-positive sample (b) and triple-negative breast cancer (TNBC) samples (c). Nine cell types in the HER2-positive samples and eight in the TNBC samples (c) were delineated using RNA expression marker genes (Supplementary Figure 1). Scale bars, 1 mm.

### Histopathologic heterogeneity is a result of diverse gene expression

Morphological heterogeneity was observed in both slides of the breast cancer samples. For instance, the HER2 positive sample exhibited a histological architecture characterized by both partial solid and partial micropapillary patterns, as well as distinct structures of both invasive ductal carcinoma (IDC) and DCIS (Fig. 2a). Transcriptomic analysis further underscored this intratumor heterogeneity, with transcriptomic variations mirroring morphological differences. We identified five clusters, including four cancer clusters and one myoepithelial cluster (Fig. 2b–d). The majority of cells in the Cancer 1 cluster corresponded to the micropapillary region. In contrast, Cancer 2 and 3 primarily aligned with solid IDC. Cancer 4 cluster, on the other hand, predominantly comprised cells that histologically structure DCIS. Finally, the last cluster enriched with myoepithelial markers was unequivocally mapped onto the histological myoepithelium. Interestingly, there were certain discrepancies between histopathological features and transcriptional clustering. For instance, certain IDC areas that were not bordered by myoepithelial cells were clustered into the Cancer 4 cluster and displayed genetic expression patterns resembling those of DCIS (Fig. 2e–f). These areas were surrounded by immune cells that separated them from other IDC regions. In contrast, there was a DCIS region exhibiting transcriptional signatures (Cancer 1 and 2) more closely resembling aggressive IDC located within the primary tumor than the adjacent DCIS (Fig. 2e).

**Fig 2.**
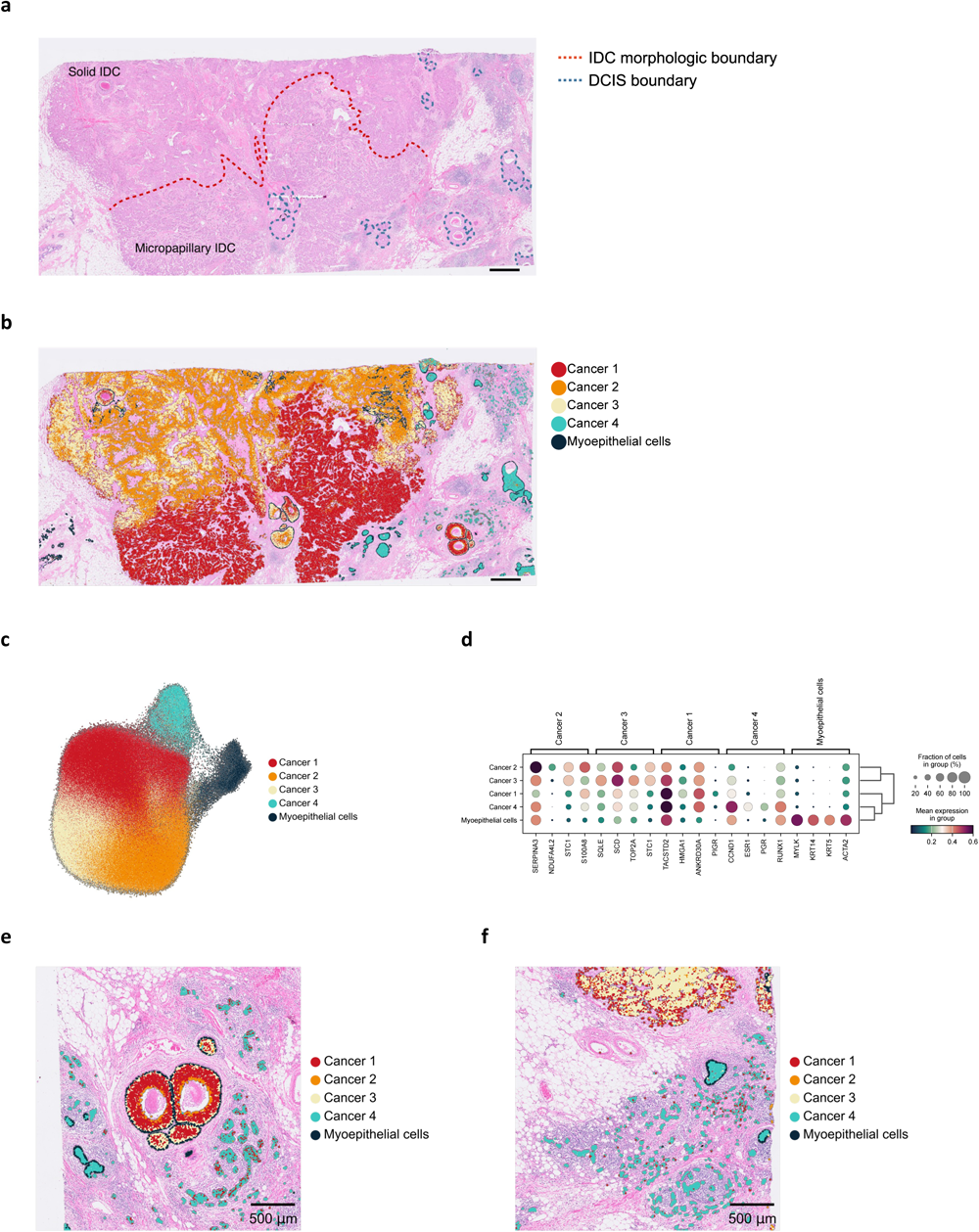
Intratumoral heterogeneity within the human epidermal growth factor receptor-2 (HER2)-positive sample. a Morphological heterogeneity annotated by a pathologist. A red dotted line separates the solid invasive ductal carcinoma (IDC) from the micropapillary IDC, while a blue dotted line aligns with the ductal carcinoma *in situ* (DCIS) area. Scale bar: 1mm. b Spatial mapping of cancer cell clusters in the HER2-positive sample, Scale bar: 1mm. c Uniform manifold approximation and projection (UMAP) embedding of cancer cells. d Matrix plot showcasing the top three genes exhibiting significant differential expression between each cluster. e–f Spatial mapping of regions with discordance between pathological terms and clusters.

In TNBC samples, we observed intratumor heterogeneity in cancer cells. Cancer cells formed two transcriptionally distinct clusters in TNBC samples (Fig. 3a, b). Cancer stage 2 was characterized by high expression of EMT genes, such as *POSTN*, *ACTA2,* and *LUM* (Fig. 3c). GSEA-supported EMT features were enriched in Cancer 2 in comparison to those in Cancer 1 (Fig. 3d). After histopathological mapping of these clusters, we observed that the genetic differences between these clusters were associated with their histopathological differences. The Cancer 1 cluster exhibited an epithelioid morphology, forming a cohesive constellation (Fig. 3e). In contrast, Cancer 2 demonstrated a mesenchyme-like morphology (Fig. 3e). Furthermore, we found evidence for the dynamics and plasticity of these two clusters. Cancer cells that underwent invasion and intravasation into the lymphovascular system exhibited concurrent EMT (Fig. 3f). Spatial trajectory analysis revealed an increase in EMT gene expression and loss of epithelial cell adhesion molecule (*EPCAM*) as the cancer cells invaded the lymphovascular area (Fig. 3g). Additionally, the expression of *CD274* and *PDCD1LG2* which potentially inhibit T cell activation, were elevated during metastasis. Interestingly, we noticed that cancer cells that had already succeeded in invading the lymphovascular space and were located away from the invasion spots exhibited morphological epithelioid features and were categorized as part of the Cancer 1 cluster and not the Cancer 2 cluster (Fig. 3h).

**Fig 3.**
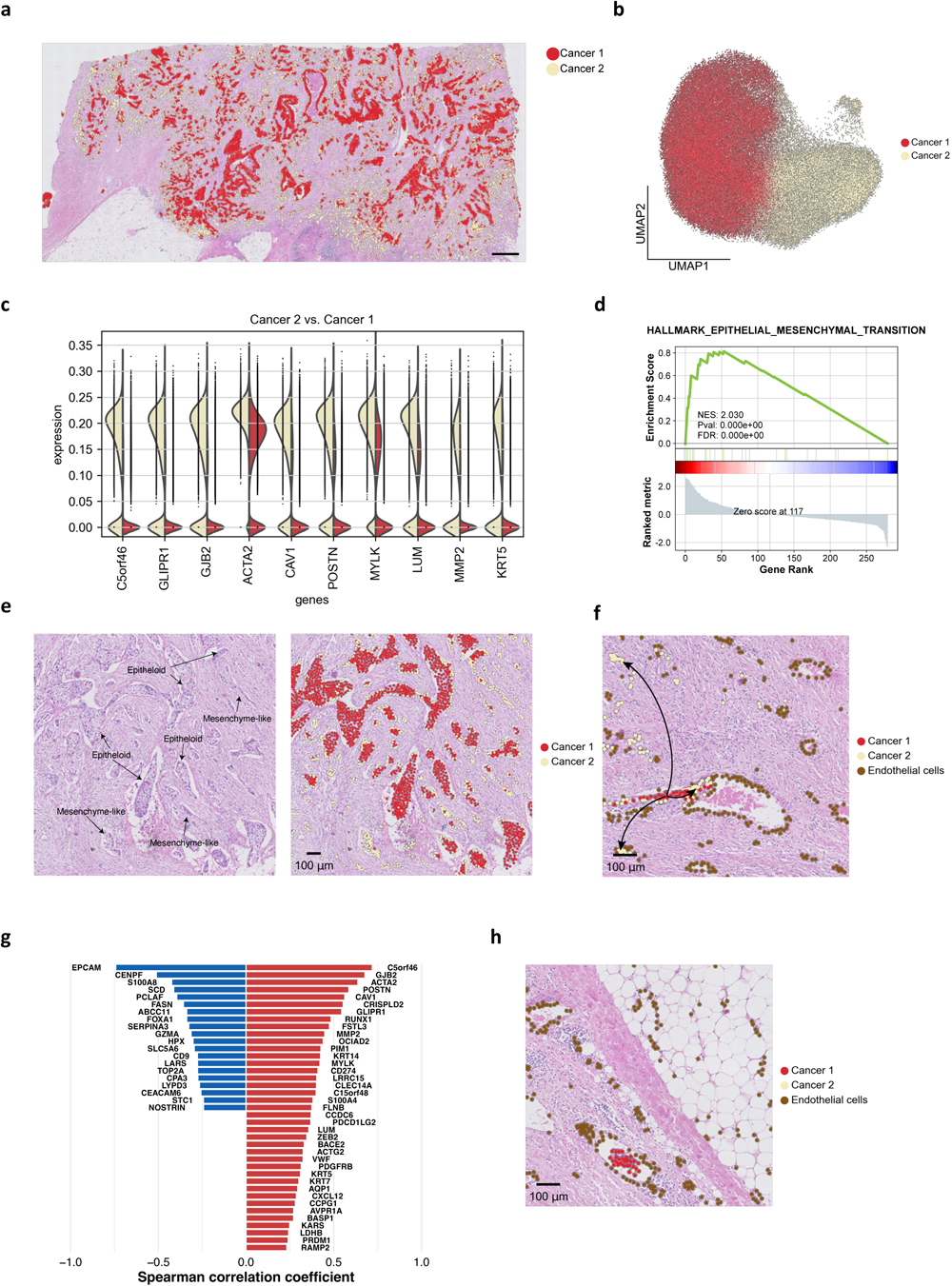
Intratumoral heterogeneity and epithelial-mesenchymal transition (EMT) process within the triple-negative breast cancer (TNBC) sample. a Spatial mapping of cancer cell clusters in the TNBC sample. b Uniform manifold approximation and projection (UMAP) embedding of the cancer cell clusters. c Split violin plots illustrating the top 10 differentially expressed genes between Cancer 1 and Cancer 2 clusters. d Gene set enrichment analysis (GSEA) of EMT pathways enriched for Cancer 2 cluster. Normalized enrichment score (NES) and *P* values are shown. e Morphologic heterogeneity aligns with gene expression-defined clusters. f Tracing the EMT process during lymphovascular invasion. g Transition markers between Cancer 1 and Cancer 2 clusters during lymphovascular invasion. h Capturing the post-invasion dynamics of cancer cells within the lymphovascular region.

### Clinical biomarkers exhibit spatial discrepancies

Next, we investigated the heterogeneity of these cancers at the genetic level. We evaluated the expression levels of the following specific markers in cancer clusters: *ESR1*, *PGR*, *ERBB2*, *EGFR*, *KRT5*, and *KRT6B* in both cancer samples. Consistent with IHC results, the HER2 positive sample exhibited strong expression of *ERBB2* (Fig. 4a), whereas the TNBC sample primarily expressed *EGFR, KRT5,* and *KRT6B*, without significant expression of *ERBB2* or hormone receptor genes (Fig. 4b). Although both samples demonstrated negative IHC results for estrogen receptor (ER) and progesterone receptor (PR), we detected mRNA expression in some cells in both samples. We also characterized the spatial gene expression patterns quantified by spatial autocorrelation using Moran’s *I* scores for each sample. Similar to the distinctive and regional morphological patterns observed in the HER2-positive sample, each marker demonstrated more clustered and regional expression patterns with strong autocorrelation in the HER2-positive samples, except *KRT6B* (Fig. 4a, b).

**Fig 4.**
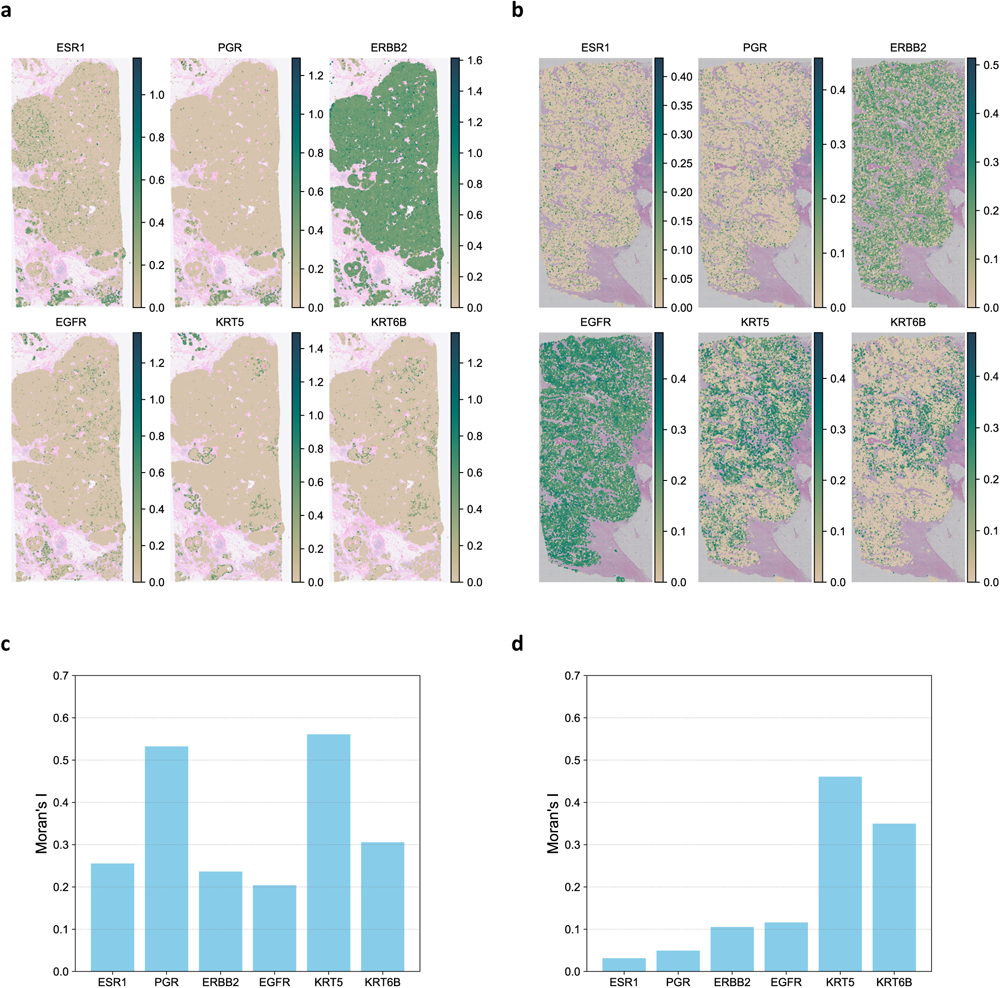
Diverse expression and spatial heterogeneity of biomarkers in breast cancer. a, b Spatial mapping of biomarker genes in HER2-positive (a) and TNBC samples (b). c, d Moran’s I global spatial autocorrelation analysis of each biomarker in HER2-positive (c) and TNBC samples (d).

### Distribution of TIME based on tumor compartments

The compartment in which TIME cells reside potentially reflects their relationship with tumor cells, other cell types, and various components within the TIME. To gain these insights, we selected a specific region with a well-defined tumor boundary and examined the proportion of TIME cells proportion in the following three spatial compartments per slide: the outer invasive margin (0 – 500 µm outside the tumor invasion front), the inner invasive margin (up to 500 µm inside), and the tumor core (Fig. 5a).

**Fig 5.**
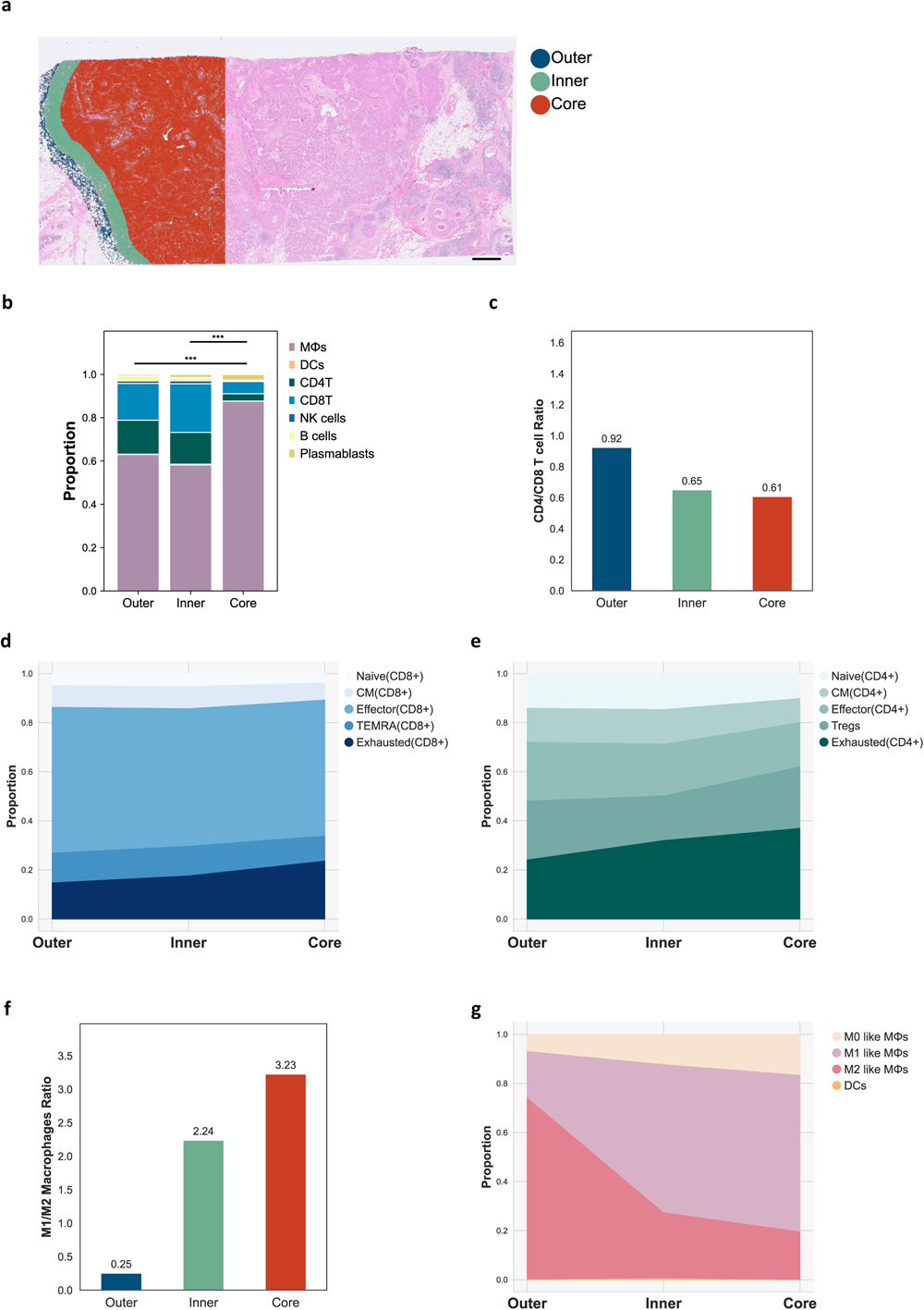
Immune landscape across different tumor compartments. a Regions of interest (ROI): outer margin, inner margin, and core. Scale bar: 1 mm. b Spatial distribution of immune cell populations. c CD4 T cells to CD8 T cells ratio, d, e Shifts in CD4 and CD8 T cell functional composition across tumor compartments. f The ratio of M1-like to M2-like macrophages, and g, The shifts in each population across tumor compartments.

In the HER2-positive sample, we observed a significant increase in the proportion of overall T cells and plasmablasts within the invasive margins (outer and inner) in comparison to the core. Conversely, the proportions of macrophages and B cells were reduced (Fig. 5b). Moreover, CD8 T cells were predominant in the core, whereas CD4 T cells were relatively abundant in the outer margin (Fig. 5c). Upon closer examination of cell types, we observed a substantial variation in cellular composition within the outer margin, inner margin, and core for both CD4 and CD8 T cells (Chi-square test: *P* < 0.001 for CD4T cells and *P* = 0.007 for CD8+ T cells). Specifically, the core exhibited a relatively reduced presence of naïve/CM T cells but a higher proportion of exhausted T cells (Fig. 5d, e). In the case of myeloid cells, M2-like macrophages were predominantly found within the invasive margin, whereas the M1-like phenotype was more prevalent in the core (Chi-square test: *P* < 0.001) (Fig. 5f, g).

We also hypothesized that T cells within the tumor core would encounter more exhausting signals, given their inability to effectively eradicate cancer cells. To investigate this, we compared the ligand-receptor pairs related to T-cell exhaustion between the inner margin and the core. Our results indicated that exhausting T cell signals were highly active in the tumor core (Fig. 6a–d).

**Fig 6.**
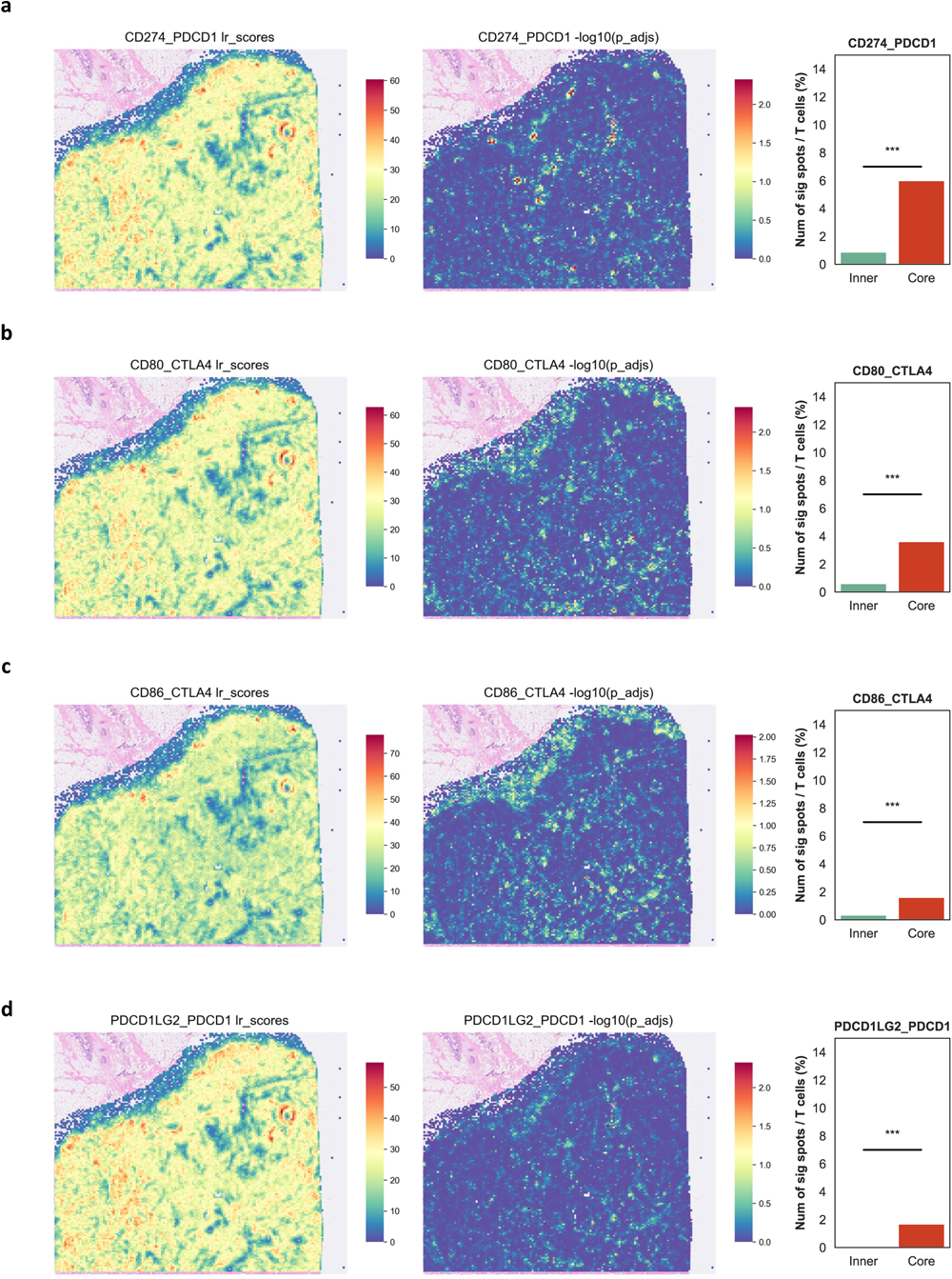
Comparative analysis of ligand-receptor pairs influencing T Cell exhaustion across compartments. a CD274-PDCD1. b CD80-CTLA4. c CD86-CTLA4. d PDCD1LG2-PDCD1.

Similar findings were observed in TNBC samples. Within the core, the number of myeloid cells surpassed the number of T cells (Supplementary Fig 5). Furthermore, the proportions of CD8 T cells and plasmablasts increased progressively from the outer margin to the core. In addition, M2-like macrophages and naïve/CM T cells were predominantly found in the invasive margin.

### Local Immune Cell Dynamics: From Training to Infiltration and Exclusion

The presence of a tertiary lymphoid structure (TLS) was detected on each slide using spatial transcriptomics. Their identification was based on their spatial cellular density and distinct histological characteristics (Fig. 7a and Supplementary Fig. 6a–d). Within these structures, a densely populated B-cell zone occupied the center and was surrounded by a peripheral region of T cells and DCs. Plasmablasts were located furthest from the center of the TLS. We also observed significant differences in the composition of each TLS, especially in CD8 T cells (Fig. 6e).

**Fig 7.**
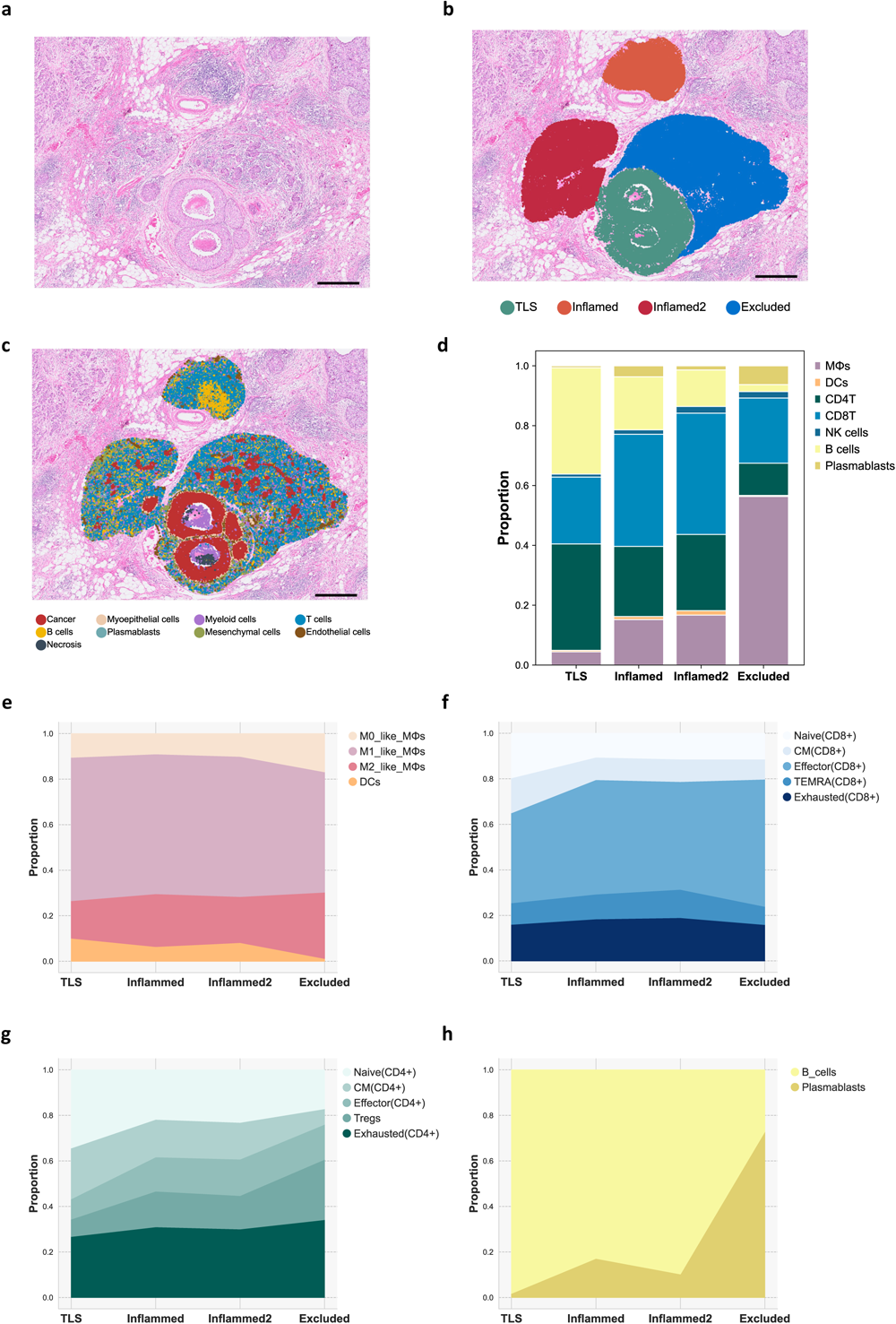
The immune landscape within the tertiary lymphoid structure (TLS) and adjacent regions in the human epidermal growth factor receptor-2 (HER2)-positive sample. a Hematoxylin and eosin (H&E) section of regions of interest (ROIs) of the TLS and surrounding areas. b Distinct regions delineated by a pathologist. c Spatial cell type mapping across these areas. Scale bar: 100 um. d Overview of the immune cell composition of these areas. – e–h Distribution of functional cell types within each immune cell cluster across the identified regions.

Comparing the cellular diversity in areas adjacent to the TLS revealed the role of TLS as a training hub for adaptive immunity. Notably, in the HER2-positive sample, we identified two tumor areas with evident immune inflammation, which were adjacent to the TLS (Fig.7a–c). Compared to these zones, the TLS exhibited a prevalent population of CD4 T cells, B cells, and DCs (Fig. 7d, e). In contrast, the immune-inflamed regions exhibited more functional T cells and plasmablasts (Fig. 7f, g). Furthermore, we identified a separate tumor region located near the inflamed area with a noticeable lack of immune cell infiltration. Analysis of this region revealed the characteristics of an immunosuppressive environment, marked by reduced CD4 T cell levels and increased frequencies of Tregs and M2-like macrophages (Fig. 7e, h).

## Discussion

Histopathology is a definitive diagnostic tool in cancer detection, largely because cancers exhibit abnormal spatial organizations within tissues.^33–35^ This distinctive tissue architecture arises from the complex interactions among various cell types. Central to these interactions is the intrinsic genetic expression inherent to each cell. By elucidating the gene expression of these individual cells within their spatial context, we can attain a genuine understanding of why cancer presents itself in such a manner and how the tumor microenvironment strategically organizes around it. Herein, we explore tissue-wide gene expression using single-cell, spatial, and *in situ* technologies to decode the spatial architecture of BC. We demonstrated that structural and morphological heterogeneities within and between tumors arise from transcriptional differences. Furthermore, we revealed that tumor microenvironments exhibit distinct compositions indexed to their spatial location within the tissues. In these locations, we observed that they engage in dynamic interactions with both tumor cells and other cell types.

The morphological heterogeneity of BC constitutes the basis for its histopathological classification.^36,37^ Furthermore, this spatial heterogeneity can be readily observed within the same tissue in daily surgical pathology. Such inter- and intratumor heterogeneities are key contributing factors for predicting treatment response and survival outcomes.^38^ Employing single-cell spatial transcriptomic technology, we delineated distinct tumor clones with both morphological and transcriptional differences. These differences range from early DCIS to advanced IDCs, capturing EMT and various histological classifications of IDC. Our results identified specific DCIS cases that could potentially evolve into IDC based on their gene expression profiles. In addition, our findings suggest that instances of myoepithelial disruption were found to be associated with co-localization of immune cells. Nevertheless, the practical implications of this co-localization of cancer cells and immune cells in these areas remains unclear. Overall, our results provide a comprehensive understanding of BC heterogeneity at the cellular level by integrating spatial information, which offers enhanced insights for interpreting histopathology, classifying BC, and predicting outcomes.

ER, PR, and HER2 expression was assessed routinely in all invasive breast carcinomas using IHC according to the recommendations of the American Society of Clinical Oncology/College of American Pathologists (ASCO/CAP)^2^. IHC methods are relatively straightforward to test but offer a limited dynamic range for quantifying marker expression compared to transcript-based methods.^39^ This limitation may restrict our ability to accurately identify the spatial distribution of each biomarker. Moreover, the lower analytical sensitivity of IHC may reduce patient eligibility for drugs targeting these markers. Our transcriptomic data exhibited distinct distribution patterns of these markers with a dynamic expression range across individual genes and slides.

Clustering plays a crucial role in single-cell data analysis. Typically, clusters determined using unsupervised methods are annotated as cell types based on the differentially expressed genes. However, high-resolution *in situ* spatial technologies often profile up to 500 genes, which account for less than 2.5% of the genome. This renders the identification of minor and functional cell types incomplete. To address these limitations, we used the well-known cell type-specific marker genes. While the near future promises accurate cell subtyping owing to technological advancements such as increments in the size of panels, our approach offers an alternative method in the interim.

Cellular organization within tissues is purposeful; specific cell types are arranged at different proximities with intent, thus enabling intercellular crosstalk and driving tissue functions, especially in patients with cancer.^40,41^ In the invasive margin, we observed a marked predominance of T cells compared to the tumor core. Notably, CD4 T cells were particularly pronounced within this T cell population; specifically, naïve/CM T cells were prominent at the invasive margin. Conversely, the tumor core was enriched with CD8 T cells, and either effector or exhausted T cells were more abundant in the core than in the invasive margin. We also demonstrated that macrophage polarization was affected by distinct tumor anatomic regions, which aligned with previous studies^42^. Collectively, our results demonstrate that immune cells *in situ* are not randomly distributed, but are influenced by their proximity to tumor cells and the tumor microenvironment.

Traditional methods for studying TLS include histology, IHC, multiplex immunofluorescence, and flow cytometry.^43–45^ While these methods can discern the spatial location and general cellular composition of TLS, they lack the resolution and depth necessary for a comprehensive understanding, as they only measure proteins and do not measure mRNAs. In contrast, single-cell spatial transcriptomic transcriptomics offers a high-resolution spatial mapping of TLSs, providing finer details about cell types and cellular organization. Notably, we observed different cellular compositions in each TLS and detected an increase in effector T cells in the inflamed regions situated close to the TLSs. These findings confirm the role of TLS as hubs for cancer-adaptive immunity beyond B cell proliferation. We also observed a unique cellular composition within the TIME, rich in Tregs and M2-like macrophages. The presented data may potentially indicate that trained immune cells from TLSs face challenges in infiltrating this cancer region owing to the prevailing TIME. The spatial, compositional, and functional characterization of a TLS is a crucial step in describing the tumor immune microenvironment at high resolution.

This study has certain limitations. Although we emphasized specific markers for minor cell subtyping, these markers are not exclusive and often overlap with each other. A clear threshold for positive expression has not been agreed upon, leading to potential bias. Furthermore, owing to the small size of the panels, it was not feasible to identify specific cell types associated with clinical outcomes among the common immune cell subtypes. Lastly, these two BC samples were from early-stage diseases that could be cured through surgical resection; therefore, we could not predict the treatment response. To date, pembrolizumab has been approved for early TNBC regardless of PD-L1 expression, and trastuzumab deruxtecan is now used in HER2 low and HER2-positive patients with BC.^5,46,47^ Therefore, this study is the first step toward revealing the predictive and prognostic biomarkers of new therapeutic strategies in BC regardless of the BC subtype.

In summary, we constructed a high-resolution spatial transcriptomic map of BC using *in situ* analysis to decipher intricate tissue architecture. Our findings enrich our understanding of intratumour heterogeneity through single-cell-level transcriptomic analysis. Furthermore, we offer enhanced insights into the TIME and demonstrate how the immune system is customized by anatomical sites, featuring distinct cell types distributed at defined ratios and spatial locations. Our research underscores the crucial role of high-resolution approaches in exploring the multifaceted heterogeneity of tumors and the TIME.

## Acknowledgments

We thank 10x Genomics for providing early technical access to the Xenium experiments.

## Author contributions

K.M. and W.Y. conceptualized and designed the study. E.S., B.R., I.W., and K.M. conducted the experiments and analyzed the data. E.S., B.L., I.W., J.Y., and W.Y. contributed to the data analysis and interpretation. E.S., B.L., J.W., and W.Y. wrote the manuscript. The corresponding authors take full responsibility for the credibility of the descriptions and data presented in this work.

## Data and Code Availability

The raw spatial transcriptomic data and codes are available at Zenodo (https://doi.org/10.5281/zenodo.10039416) and our analysis code have been uploaded to github at https://github.com/SGIlabes/BRCA_xenium.

