## Supplementary Figure for "Decoding spatial organization maps and context-specific landscapes of breast cancer and its microenvironment via high-resolution spatial transcriptomic analysis"

### Supplementary Fig. 1

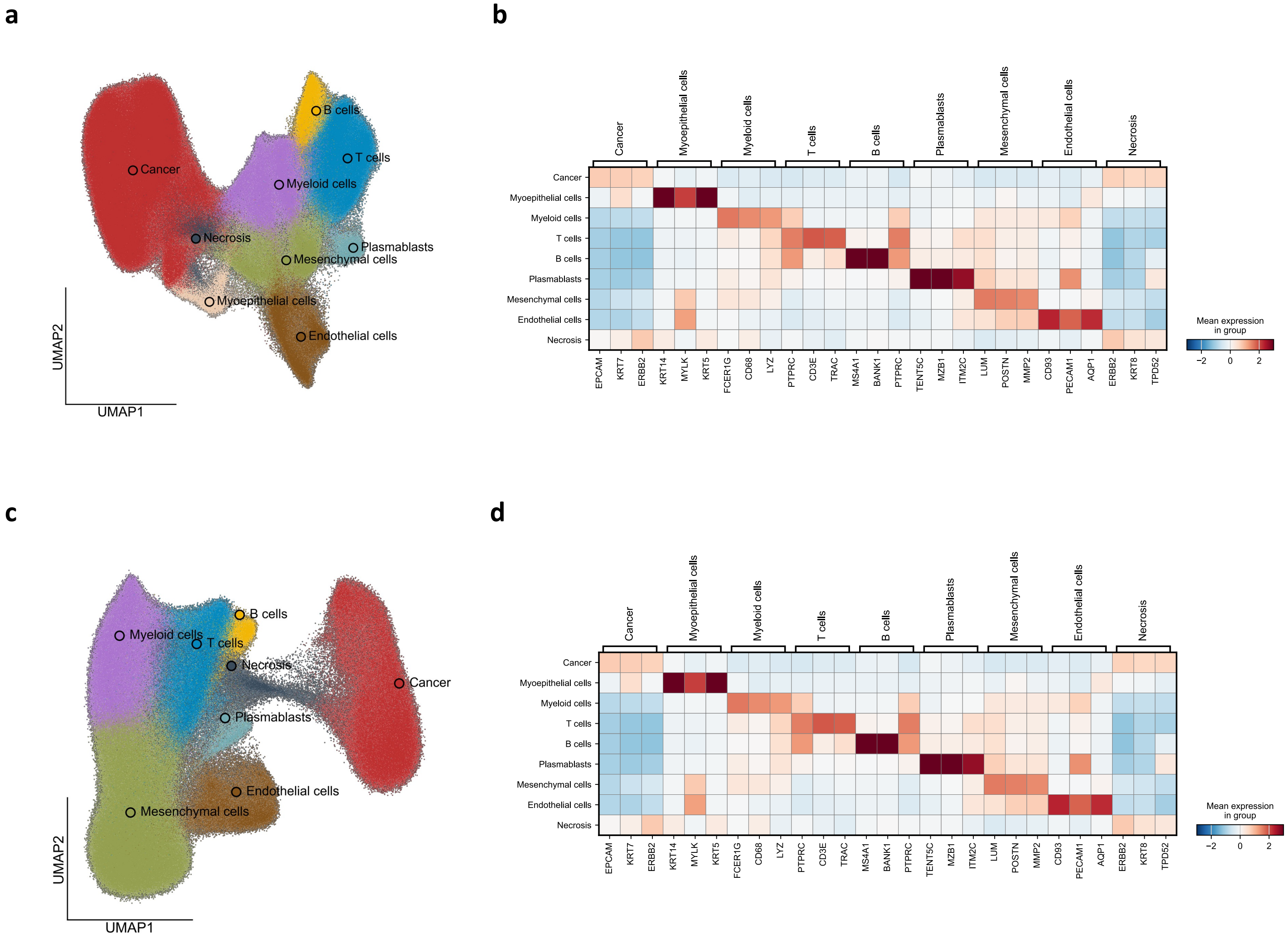

### Supplementary Fig. 2

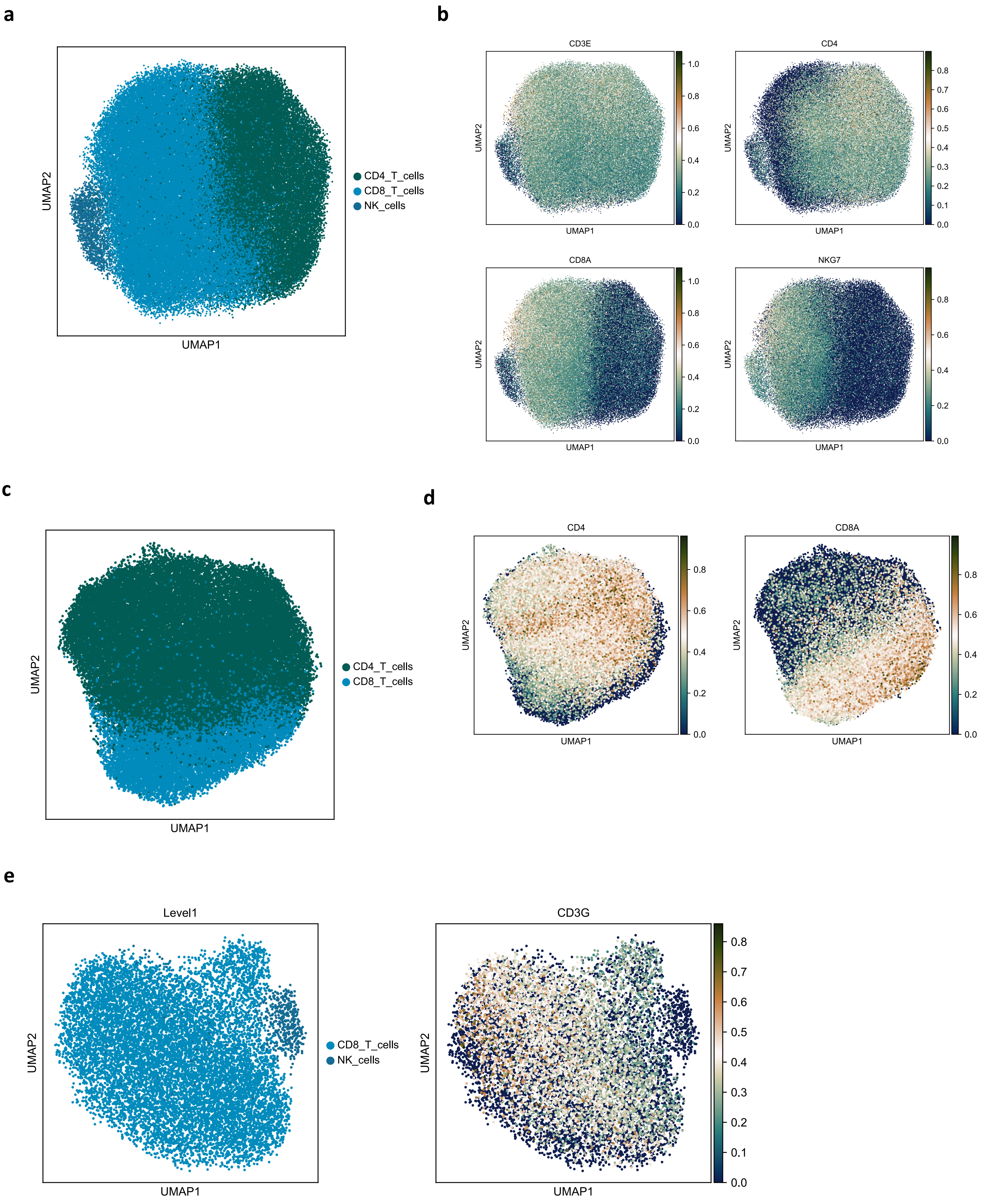

#### Supplementary Fig. 3

**a**

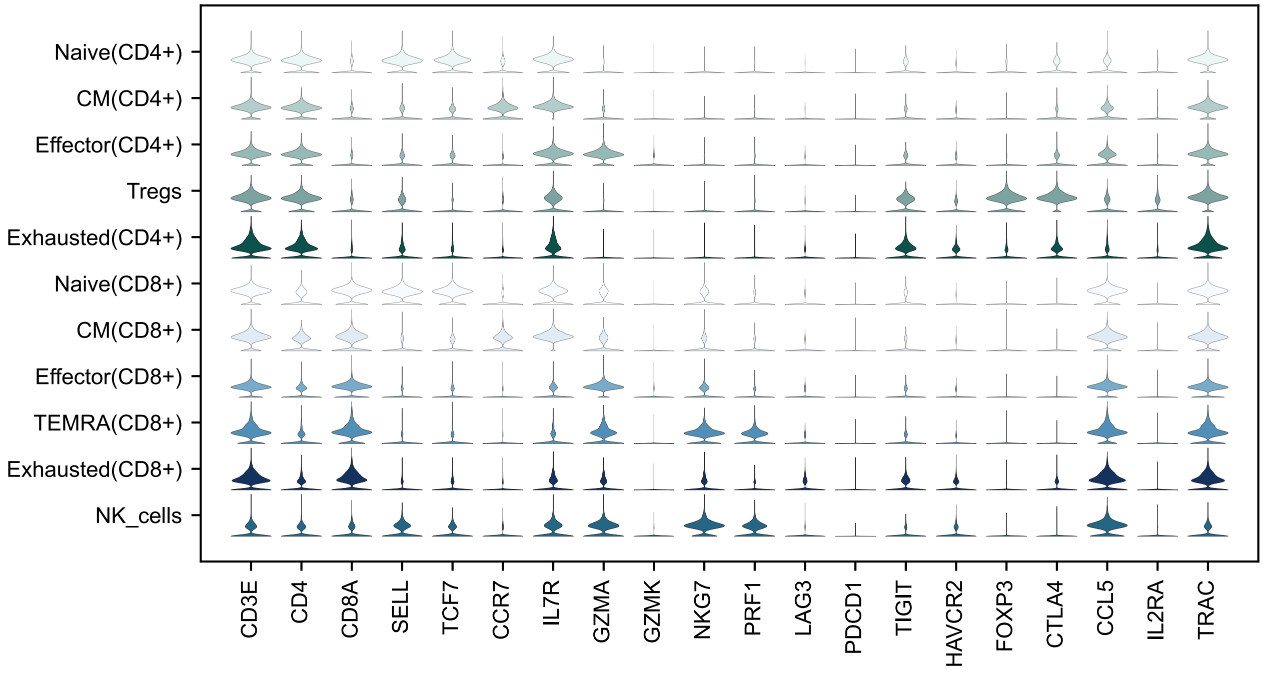


**b**

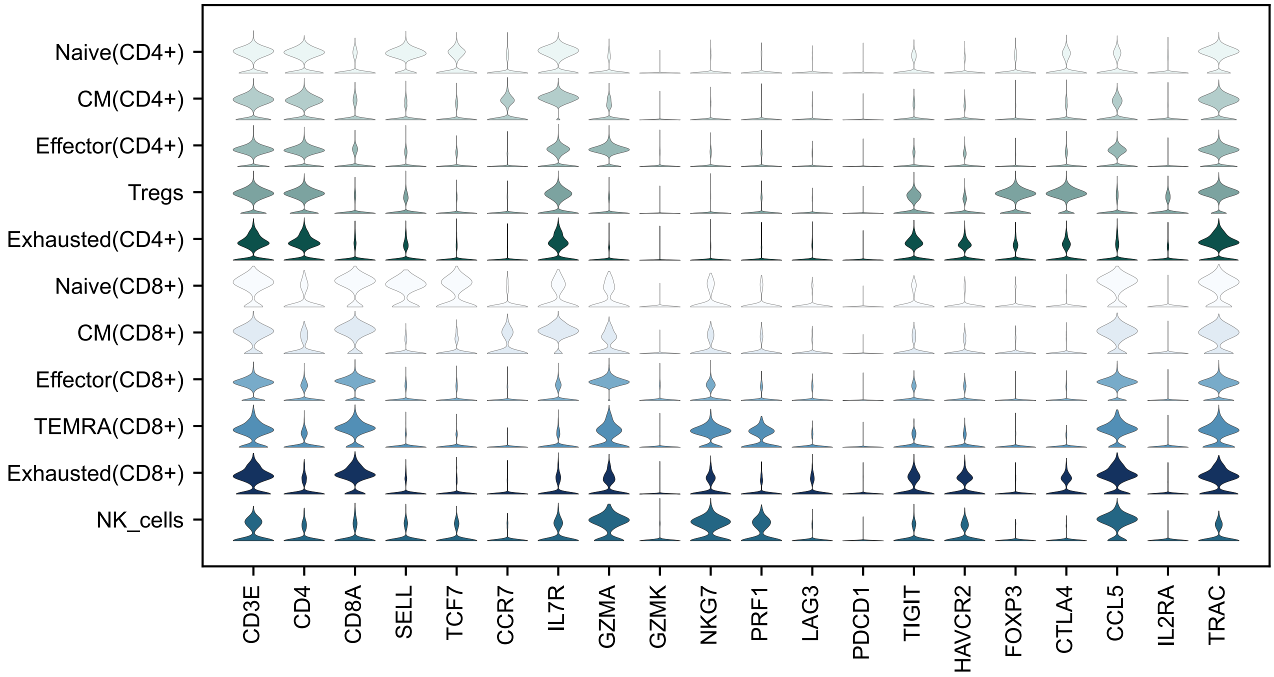

#### Supplementary Fig. 4

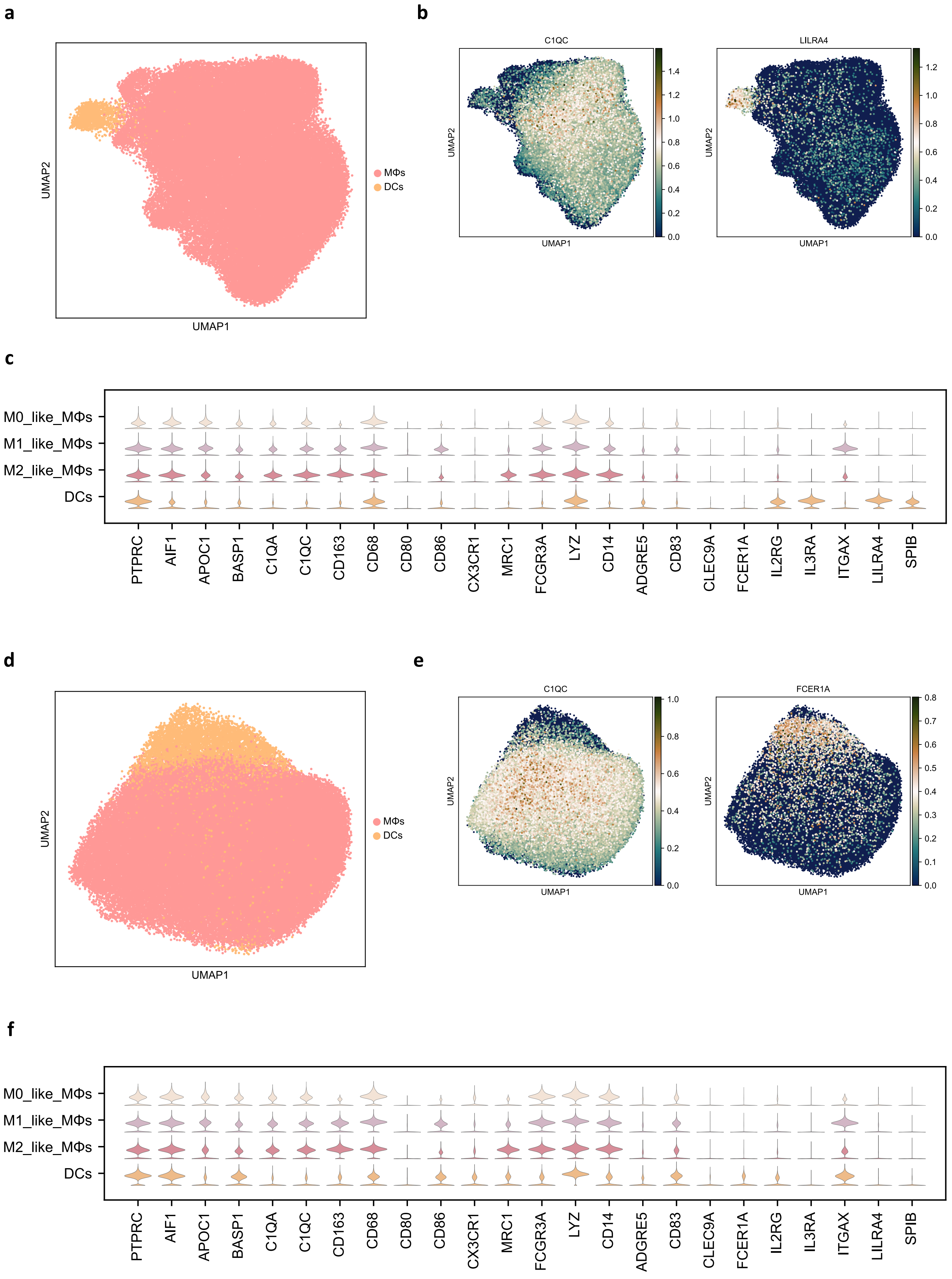

### Supplementary Fig. 5

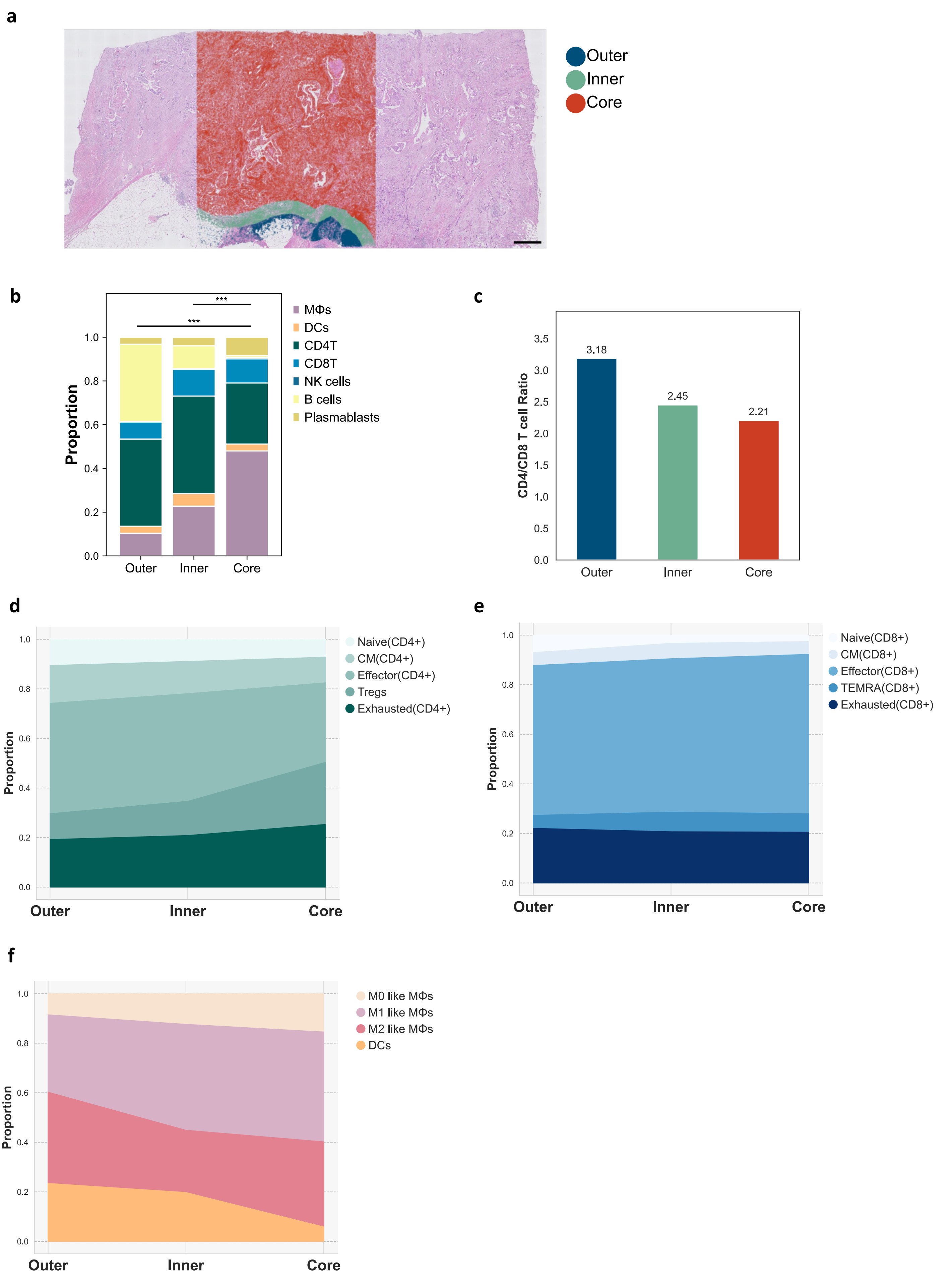

### Supplementary Fig. 6

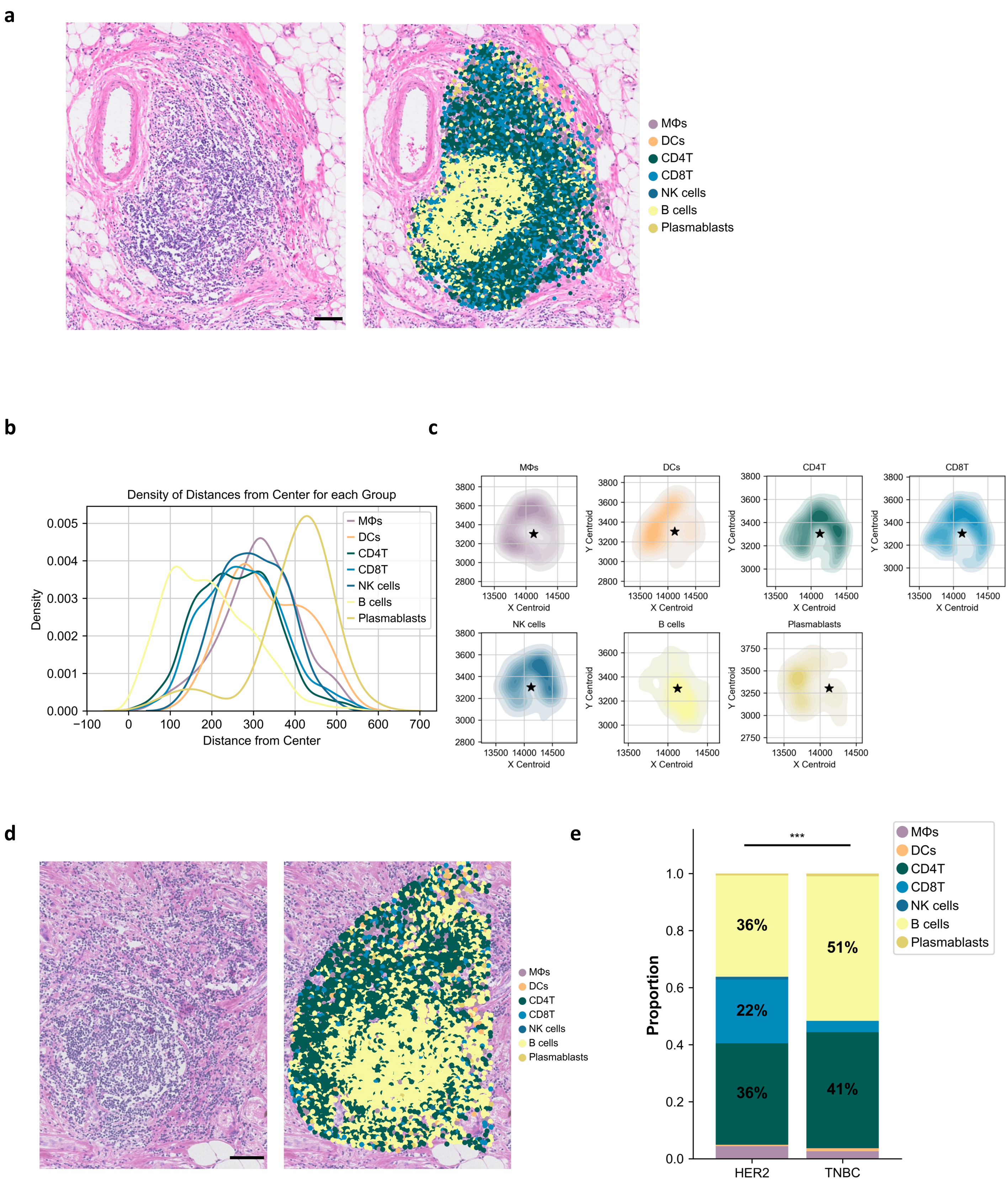
